# Patellar tendon-tap intensity modulates response gain but not variability across cortical proprioceptive and stretch-reflex responses

**DOI:** 10.64898/2026.09.21.753125

**Authors:** Feiyue Li, Anni Byman, Junru Chen, Toni Mujunen, Harri Piitulainen

## Abstract

Cortical proprioceptive processing to passive movements can be quantified with evoked and induced responses in magnetoencephalographic (MEG) signals. We examined how intensity of proprioceptive stimulation (i.e., tendon-tap evoked movement) scales cortical processing of proprioceptive afference from the knee joint, and whether it affects stimulus-to-stimulus response variability.

Twenty-one healthy volunteers (28.2±5 yr, 10 females) underwent a total of 100 right patellar tendon taps (5.0−6.0s inter-stimulus interval) evoked by a novel MEG-compatible stimulator under *higher* and *lower* intensity conditions. Strength of cortical evoked and induced MEG and electromyographic responses from the vastus lateralis and medialis muscles were quantified, and their variability were assessed using a matched sliding-window coefficient of variation (CoV).

Higher intensity stimulation elicited significantly stronger cortical evoked (∼11%) and induced (∼17% for beta suppression and ∼18% for beta rebound) responses, and muscular response (∼27%) than lower intensity stimulation (p<0.05). The stimulus intensity did not affect response variability, with comparable CoV values between higher and lower intensity condition (p>0.05), but the muscular responses showed greater variability than the cortical ones in both intensity conditions (CoV: ∼30% vs. ∼10%, p<0.001). In addition, the variability in cortical and muscular responses were not significantly correlated (p>0.05).

Our results indicate that patellar tendon-tap intensity scaled cortical and muscular response gain, but did not measurably alter response variability. The distinct variability profiles of cortical evoked fields and stretch-reflex muscular response suggest that proprioceptive afference evoked by tendon-tap is transformed differently across cortical and spinal levels.

## Introduction

Proprioception is the sense of the body position and movement in space (Dietz, 2002; Proske and Gandevia, 2012), and it plays a fundamental role in movement control, postural regulation and the continuous integration of sensory feedback with motor commands (Han et al., 2015; Kavounoudias et al., 1999; Kiehn, 2016). The primary source of proprioceptive information arises from muscle spindles (Proske and Gandevia, 2012; Tuthill and Azim, 2018), which provide information about changes in muscle length during movement to the central nervous system. In humans, proprioceptive stimulation has been shown to elicit robust cortical responses in the primary sensorimotor (SM1) cortex, and these responses can be measured non-invasively using magnetic resonance imaging (MRI; Nurmi et al., 2018; Nurmi et al., 2021), electroencephalography (EEG; Giangrande et al., 2024; Piitulainen et al., 2020; Qiu et al., 2016) and magnetoencephalography (MEG; Alary et al., 2002; Chen et al., 2025; Lange et al., 2001; Mujunen et al., 2022; Piitulainen et al., 2015). These responses open a window to quantify the features of cortical proprioceptive processing.

Previous studies suggest that cortical responses are sensitive to the properties of somatosensory peripheral electrical stimulus, including stimulus intensity. With the median nerve electrical stimulation, EEG and MEG studies have showed that increasing stimulus intensity generally enhances early cortical responses, indicating that stronger afferent input can increase cortical response strength, although this scaling may become non-linear or plateau at higher intensities (Gobbele et al., 2008; Hewitt et al., 2022; Jousmaki and Forss, 1998; Smith et al., 2003; Torquati et al., 2002). Insausti-Delgado et al. (2020) further showed that graded electrical stimulation progressively increased beta event-related desynchronization strength over the sensorimotor cortex, supporting an intensity-dependent modulation of beta rhythm. By contrast, evidence from mechanically evoked proprioceptive stimulation is less consistent, probably because different mechanical paradigms generate distinct afferent patterns. In passive finger movement paradigms, for example, cortical response strength has been reported to remain invariant across movement ranges in MEG (Nurmi et al., 2023), whereas higher movement frequency can intensify fMRI-BOLD signal associated with the cortical proprioceptive processing (Nurmi et al., 2018). Simultaneous proprioceptive stimulation of multiple fingers of the same hand can also enhance corticokinematic coherence strength (CKC) in MEG when compared to stimulation of one finger (Hakonen et al., 2022). Thus, cortical response may depend not only on mechanical stimulus strength, but also on how widely the stimulus-related afferents are activated. Similarly, Zhao et al. (2025) investigated the relationship between CKC strength and passive movement parameters in the lower limb (i.e., hip, knee and ankle joints) by using EEG. They showed that CKC strength increased progressively with movement range at low movement frequencies, whereas this range-dependent increase was not consistently observed at higher movement frequencies.

It is still unclear whether stimulus intensity systematically scales cortical proprioceptive processing during naturalistic lower limb mechanical stimulation. Most of previous evidence has focused either on electrical stimulation of peripheral nerves or on passive movements of the hand, whereas less is known from lower limb stimulation of the knee joint. Tendon tap provides a distinct form of mechanical stimulation: unlike passive movement, it produces a brief, dynamic stretch of the muscle-tendon unit and elicits a spinal stretch reflex response. An early EEG study showed that mechanical stimulation, tendon taps, evoked cortical somatosensory responses in accordance with those elicited by peripheral median nerve stimulation (Larsson and Prevec, 1970). More recent tendon-tap studies have further shown that stronger mechanical tap increases amplitude of the muscular patellar tendon reflex response (Chandrasekhar et al., 2013; Tham et al., 2013; Tsuji et al., 2021). Patellar tendon tap provides a naturalistic way for examining proprioceptive afference from the knee extensors at both spinal and cortical levels, analogous to passive finger or ankle movements evoked by movement actuators to stimulate the peripheral proprioceptors in the previous studies (Piitulainen et al., 2015; Piitulainen et al., 2018; Walker et al., 2020).

Cortical proprioceptive processing can be characterized using both stimulus-locked evoked response and induced beta band modulation measured with MEG or EEG (Alary et al., 2002; Druschky et al., 2003; Giangrande et al., 2024; Illman et al., 2022; Mujunen et al., 2025; Nurmi et al., 2023). Evoked response primarily reflects the cortical activation in time, phase locked to proprioceptive stimulation, whereas the induced response reflects the cortical excitation-inhibition dynamics of the sensorimotor system following afferent input (David et al., 2006; Kilavik et al., 2013). Specifically, the amplitude of beta band power in the SM1 cortex is reduced shortly after stimulus (suppression, or event-related desynchronization, ERD), followed by an increase in the amplitude (rebound, or event-related synchronization, ERS). The beta suppression reflects an activation, and the rebound represents a deactivation or inhibition of the cortex (Barone and Rossiter, 2021; Neuper et al., 2006; Takemi et al., 2013). The evoked and induced responses are typically strong and observable in most individuals (Giangrande et al., 2024; Mujunen et al., 2022; Toledo et al., 2016; Walker et al., 2020). Moreover, our previous study has revealed that they are reproducible at the group level for MEG in the lower limb (Mujunen et al., 2022).

In this study, we primarily aimed to examine whether cortical processing of proprioceptive afference is amplified by stronger mechanical patellar tendon tapping. Based on the evidence reviewed above, we hypothesized that higher intensity stimulation would result in stronger cortical responses (stronger evoked field and beta power modulation), as well as stronger reflex-related muscular responses. Our secondary aim was to clarify whether cortical response variability is coupled with muscular response variability. We further hypothesized that the response variability would be lower at cortical than spinal level, since cortical responses reflect more integrated population-level processing of peripheral afference (Groh et al., 2014; Proske and Gandevia, 2012), whereas the spinal cord generates the rapid response to the stimulation and is more directly affected by trial-to-trial fluctuations (Macefield and Knellwolf, 2018; McNeil et al., 2013; Reschechtko and Pruszynski, 2020). To test these hypotheses, we used *higher* and *lower* intensity patellar tendon taps to elicit the reflex and stimulate the proprioceptors in the knee extensors during MEG and electromyographic (EMG) recordings.

## Materials and methods

### Subjects

We recruited twenty-three healthy participants with no history of movement disorders or neuropsychiatric disease (11 females, mean ± SD, age = 28.3 ± 5 years). According to the Waterloo footedness test (Elias et al., 1998), the inventory score was 8 ± 4 on a scale from −20 to 20, indicating that the predominance of right-leg volunteers (22 out of 23 participants). All participants received an explanation of the whole protocol before signing the informed consent. The study had prior approval from the University of Jyväskylä Ethics Committee in accordance with Declaration of Helsinki (approval number: 694/13.00.04.00/2023).

### Experimental design

Figure 1 shows the participant with the novel MEG-compatible stimulator that evoked the patellar tendon reflex to generate a rapid contraction of the quadriceps muscles in the knee joint. This novel stimulator consisted of a supporting PVC-plastic frame and a hammer. The hammer was first manually held to the initial position when the visual cue was provided thus releasing the hammer, and then allowing it to fall in a gravity-driven arc toward the patellar tendon. The hammer was moved back to the initial position after each first contact with the patellar tendon. For further information on the novel stimulator, see our recent work (Li et al., 2026). Depending on the hammer release height, there were two stimuli conditions: higher intensity and lower intensity (Figure 1B). The order of the intensity condition was randomized for the first participant and then alternated for the following participants, i.e., if higher intensity was firstly delivered for the first participant, then the next participant started with lower intensity. A total 100 of the patellar tendon tapping with an inter-stimulus interval of 5.0−6.0s were delivered for each condition. The stimulus timing was indicated to the experimenter through light signal by optical fiber using Presentation software (Ver. 21.1, Neurobehavioral Systems Inc., Albany, CA, USA).

**Figure 1.**
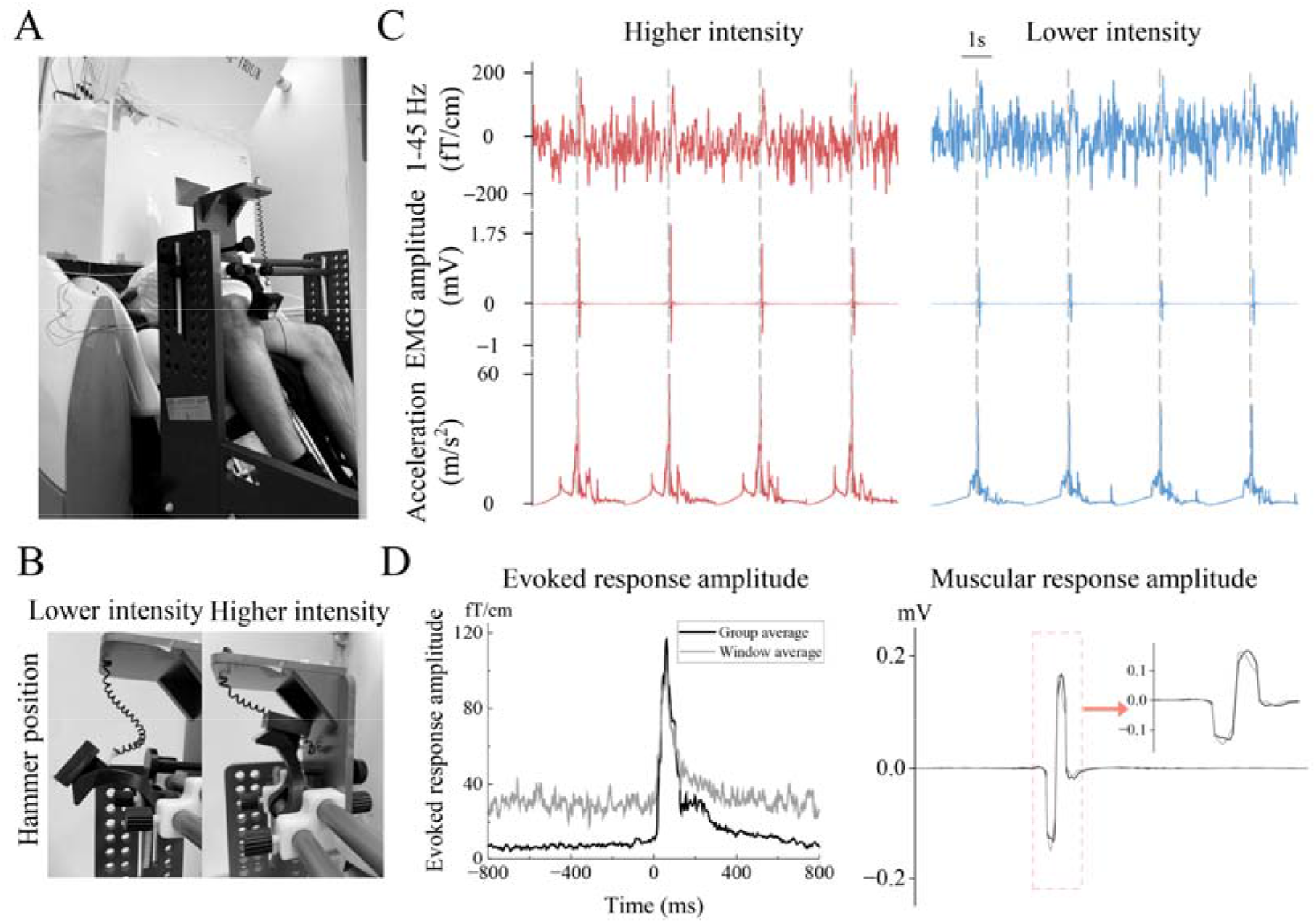
Experimental protocol and signals. (A) Participant’s right leg placed with knee joint at 90°. The hammer head contacted the patellar tendon perpendicular (90°) with the stimulation site. (B) The hammer initial dropping position for lower (left) and higher (right) intensity conditions. (C) Continuous 1−45 Hz MEG from the peak gradiometer channel, EMG from the vastus lateralis (VL) and acceleration signals from one representative participant in both intensity conditions (red, higher intensity and blue, lower intensity). Dashed vertical lines indicate stimulus onset (i.e., the tendon tapping). (D) Example of cortical evoked field (left) and muscular response (right) averaged with 4 stimuli (one window average, gray line) and all ∼100 stimuli (full average, black line) in higher intensity condition.

During MEG recording, the participant was sitting with their foot of the stimulated right leg on a pedal-like support of the custom-made MEG chair embedded with force transducers (University of Jyväskylä, Finland), that was used to record the reflex-related knee-extension force. The stimulator was initially positioned in front of the MEG chair and aligned with the reference stimulation point on the patellar tendon identified during the preparation. The guide rods and the horizontal position of the hammer were adjusted to ensure that the hammer head contacted the patellar tendon perpendicular to the tendon surface at the reference stimulation point. In addition, the stimulator position was then fine-tuned by delivering several test taps and monitoring the reflex response. If the initial reference point did not elicit a clear and reproducible reflex, the stimulator and hammer were shifted slightly along the patellar tendon until a prominent quadriceps response was obtained. The final position was then kept constant throughout the MEG recording. Foam padding and adjustable straps were used around the right thigh and lower shin to ensure comfort and to minimize head movements that may cause motion artefacts in MEG signals. The participant’s arms were resting on a pillow on their lap. To further prevent head movement inside the MEG helmet, pieces of foam cushion with the net cap were placed on both sides of the head, between the temporal sides of the head and the MEG helmet. The light acoustic noise from the hammer was masked with Brownian played from flat panel speakers inside the shielded room. In addition, visual contact to the hammer and the knee joint was blocked with a paperboard while allowing the participant to fixate their gaze on a fixation cross placed on the fluorescent screen in front of the participant.

### Signal acquisition

### MEG

MEG signals were recorded with a 306-channel whole-scalp neuromagnetometer system (Elekta Neuromag TRIUXTM, Elekta Oy, Helsinki, Finland). Signals were sampled at 1000 Hz with 0.1−330 Hz passband. In addition, eye blinks were collected with two electro-oculography and the ground electrode was placed on top of the right clavicle. During preparation, five head position indicator (HPI) coils were attached (three on the forehead and one above each ear lobe) to define and follow the participant’s head position with respect to the MEG sensors. The location of the HPI coils, three anatomical landmarks above nasion and two preauricular points, and 200 additional points from the scalp surface were determined using a 3-D digitizer (Isotrak, Polhemus, Colchester, VT, USA). The head position with respect to the sensor array was checked at the beginning of each measurement session and was recorded during the measurement.

### Kinematics

The hammer acceleration was collected with a 3-axis accelerometer (ADXL335 iMEMS Accelerometer, Analog Devices Inc. Norwood, MA, USA) placed on the top head of the hammer. The peak acceleration magnitude (i.e. Euclidean norm of three acceleration axis) and its latency were calculated to determine the stimulus onset. Acceleration signal was bandpass filtered from 0.1−330 Hz with the sample frequency of 1000 Hz, time-locked to MEG signals.

### EMG and reflex force

The muscular responses to the patellar tendon reflex were recorded from the vastus lateralis (VL) and vastus medialis (VM) muscles using bipolar EMG with two pairs of Ag/AgCl electrodes (Ambu Neuroline 720 15-K/C/12, Denmark). The electrode placements were based on the study by Barbero et al. (2012) aiming to avoid placement on the main innervation zone(s). Before attaching the EMG electrodes, body hair and dead skin were removed with razor and sandpaper respectively. The knee reflex force was recorded with a load cell embedded in the MEG chair. The quality of EMG and reflex force signals were confirmed before onset of the actual recordings, and bandpass filtered from 0.1−330 Hz with the sample frequency of 1000 Hz, time-locked to MEG signals.

### Data preprocessing and analysis

First, to find event timing in MEG and EMG data, the acceleration signal was used to determine stimulus onset (i.e., the tendon tapping). The acceleration signal was bandpass filtered at 1−195 Hz and the Euclidian norm of the three orthogonal signals (i.e., acceleration magnitude signal) was calculated and combined as ‘peak magnitude’. Stimulus onset was defined as the timepoint corresponding to the peak magnitude separated for each stimulus, participant and both intensity conditions.

### MEG preprocessing

Custom-made Python script was used to manually identify the noisy MEG sensors in MEG signals for each participant. The averaged head coordinates, calculated from the head positions in different stimulation conditions were used as reference head position in the Maxfilter software (v3.0; Elekta Oy, Helsinki, Finland) for head movement compensation, with the signal-space separation method with temporal extension (tSSS) to reduce external interference and head movement effects. Then, MEG signals were decomposed into 30 components through the independent component analysis function to remove the components related to eye blinks and muscular activity and filtered between 1−40 Hz using a zero-phase finite impulse response filter with MNE-Python (Gramfort et al., 2013).

### EMG and force amplitudes of muscular response

First, EMG signals were bandpass filtered at 10−295 Hz and epoched from −2000 to 2000 ms with respect to stimulus onset. Then, the first epoch was discarded to minimize the effect of early tapping reaction, and the remaining epochs were averaged. To quantify muscular response during higher and lower intensity conditions, the peak-to-peak amplitude of EMG data was computed for VL and VM muscles. Similar processing was done for the force signal, but with bandpass filter at 1−80 Hz. The peak-to-peak force responses were averaged separately for higher and lower intensity conditions.

### Evoked field

The preprocessed MEG signals were filtered with band frequency 1−45 Hz and epoched from −2000 to 2000 ms with respect to stimulus onset. Epochs that contained the signal amplitude exceeding amplitude of 4pT for magnetometers and 4pT/cm for gradiometers were excluded from the analysis to avoid possible noisy segments in the averaging. Then, epochs were averaged separately for each channel and condition. The magnetometer signals were replaced by combining the vector sum of two gradiometer signals for each pair. The gradiometer pair with peak activation in the parietal area was chosen for the final analysis. The peak gradiometer pair was selected independently in each intensity condition.

### Beta rhythm modulation

The induced response (i.e., ∼20 Hz beta rhythm modulation) was quantified using the temporal spectral evolution (TSE) method in both intensity conditions. The preprocessed MEG signals were bandpass filtered at 12−30 Hz and epoched from −2000 to 2000 ms with respect to stimulus onset. After epoching, the evoked field was subtracted from the data (David et al., 2006), which was rectified and averaged time-locked to stimulus onset. As with the evoked field, the gradiometer showing the peak amplitude of beta suppression and rebound was selected from the parietal area for each intensity condition and participant separately. If there was no clear and visible beta modulation that exceeded three standard deviations of the baseline signal amplitude with the period from −1800 to −100 ms, the participant was excluded from the final analysis. If the strongest rebound and suppression were on different channels, the respective peak channels were selected separately for further analysis. To quantify the amplitude of beta suppression and rebound, the baseline correction (from −1800 to −100 ms) was applied to the averaged data and the peak values were converted to the relative value (in percentage) with respect to the baseline.

### Sliding-window estimation of response variability

The single-trial MEG signal typically exhibits a relatively low signal-to-noise ratio, and thus individual trials estimation is difficult even in case of the most prominent MEG responses. Therefore, we used a sliding-window coefficient of variation (CoV) analysis to estimate response variability from short averages of consecutive stimuli and applied the same windowing procedure to both cortical and muscular responses. For cortical response, the signal was extracted from the same peak gradiometer pair (i.e. their vector sum) as in the evoked field analysis (see above). Four consecutive stimuli were used per window, since this window preserved an identifiable evoked field waveform while providing multiple window-level estimates within each intensity condition (see Fig.1D).

Within each participant and condition, the MEG or EMG data was divided into 50% overlapping windows of four stimuli with a step size of two stimuli. The first window was excluded to minimize the effect of startle or orienting responses when tendon tap was first introduced at the beginning of each condition. Then, the stimuli within each window were first averaged, and response strength was quantified as the root mean square (RMS) of the averaged waveform from 0 to 100 ms with respect to stimulus onset (i.e. the tendon tapping). Finally, response variability was calculated as the CoV of RMS across all 4-stimulus windows, defined as the standard deviation divided by the mean (CoV = SD/mean).

### Statistical analysis

All statistical tests were conducted in IBM SPSS software (v. 27.0). Shapiro-Wilk test was used to assess the normality of the data. One-way repeated measures multivariate analysis of variance (MANOVA) was carried out to test whether the stimulus intensity affects cortical and muscular responses with normally distributed variables (peak evoked amplitude, peak beta suppression, peak beta power latencies and CoV of VL and VM muscles). In the case of statistically significant main effects, the paired t-test was used to determine differences between higher intensity and lower intensity conditions. In addition, a non-parametric Wilcoxon signed rand test was conducted to analyze significant differences in non-normally distributed parameters, including reflex force, peak acceleration magnitude, peak muscular response, peak evoked fields latency, peak beta rebound and CoV of evoked fields. Lastly, the Benjamini-Hochberg false discovery rate correction (FDR) was used to adjust the significant level (i.e., p value) for multiple comparisons. Spearman correlation was used to calculate correlations between cortical and muscular response variability.

## Results

We only included participants with robust cortical and muscular responses in both higher and lower intensity conditions to the final analysis (n = 21, 10 females, age = 28.2 ± 5 years). We first confirmed the stretch-reflex response from the EMG recordings; participants without a clear muscular response were excluded before cortical analysis. We then inspected cortical evoked and induced responses to confirm identifiable stimulus-locked activity in both intensity conditions. Based on this criteria, two participants were excluded from the final analysis: one because of the lack of clear muscular response and induced response in both intensity conditions, and one due to unidentifiable induced response in higher intensity condition. After preprocessing, only 5% of stimulus epochs in muscular response were excluded in further analysis, 2% in evoked field and 3% in induced response for higher intensity condition, and 3%, 1% and 3% for lower intensity condition, respectively. Figure 1C shows pre-processed signals in one representative participant for both higher and lower intensity conditions.

### The strength of proprioceptive stimulus

Figure 2 top panels show the results for stimulus kinematics in higher and lower intensity conditions. Figure 2A left panel illustrates group averaged time-series for stimulus kinematics. As expected, Wilcoxon test showed that peak acceleration magnitude was significantly higher in higher intensity than in lower intensity condition (W = 8, p = 0.002). Table 1 shows stimulus kinematics.

**Figure 2.**
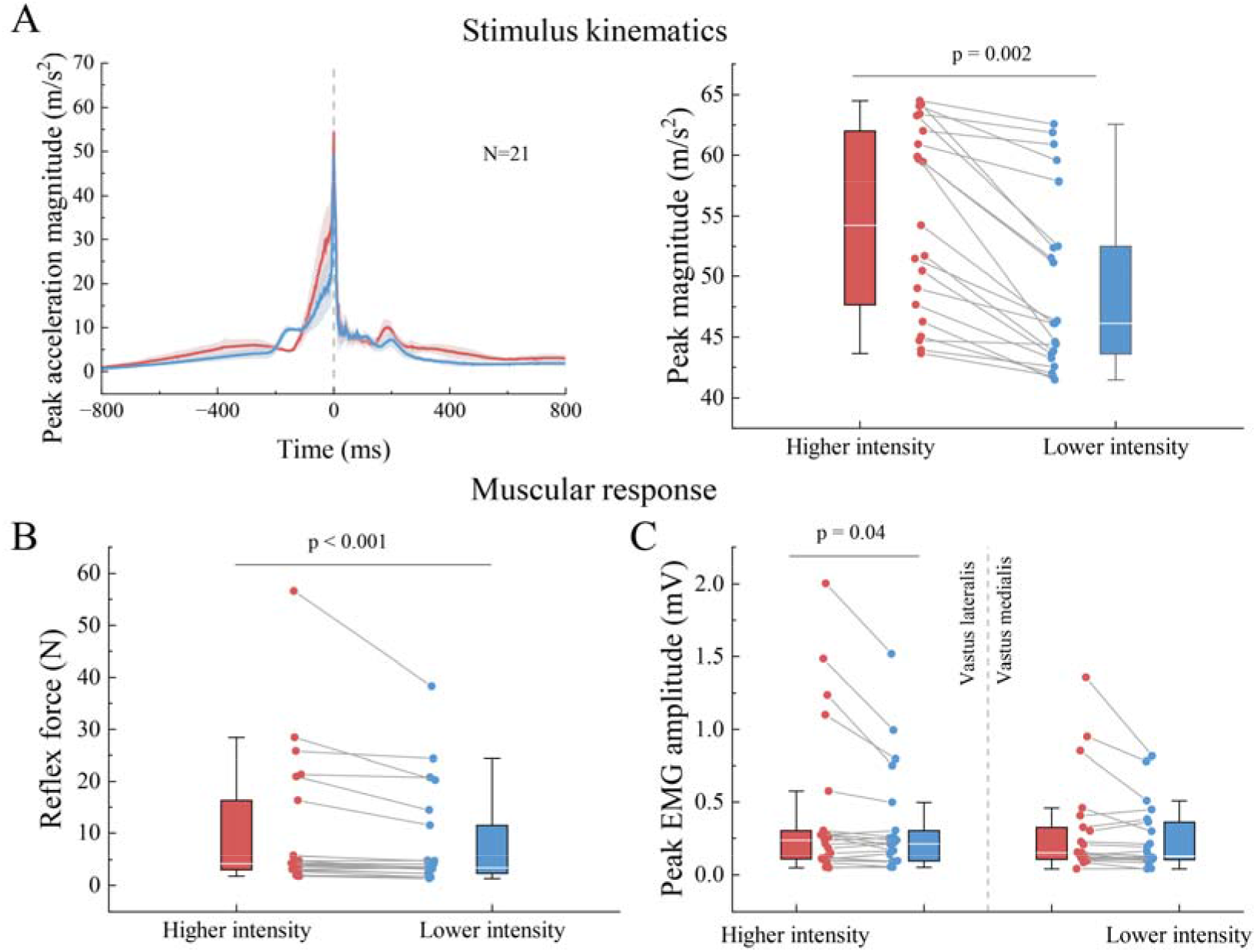
Stimulus kinematics, reflex force and muscular response in both higher and lower intensity conditions. (A) The left panel shows grand average acceleration magnitude across participants for higher (red) and lower (blue) intensity conditions. The comparison of peak acceleration magnitude for both higher and lower intensity conditions (right panel). Initial peak value (vertical grey dashed line) was used to determine stimulus onset. (B) Boxplots of reflex force for both intensity conditions. (C) Peak values as boxplots for VL and VM EMG amplitude for both intensity conditions. Individual values are shown with points.

**Table 1.** Pooled kinematics, reflex force, muscular, evoked and induced response parameters (mean ± SD) for higher and lower intensity conditions.

|  | Higher intensity | Lower intensity | p-value |
| --- | --- | --- | --- |
| **Stimulus kinematics** |  |  |  |
| Peak magnitude ($\text{m/s}^2$) | $54.58 \pm 7.68$ | $49.49 \pm 7.50$ | **0.002** |
| **Muscle response** |  |  |  |
| Reflex force (N) | $10.5 \pm 13.6$ | $8.3 \pm 9.9$ | **< 0.001** |
| VL EMG amplitude (mV) | $0.44 \pm 0.54$ | $0.34 \pm 0.38$ | **0.04** |
| VM EMG amplitude (mV) | $0.31 \pm 0.34$ | $0.25 \pm 0.23$ | 0.15 |
| **Evoked fields** |  |  |  |
| Peak amplitude (fT/cm) | 149.15 ± 55.92 | 133.79 ± 45.24 | **0.03** |
| Latency (ms) | 57 ± 9 | 59 ± 9 | 0.53 |
| **Induced responses** |  |  |  |
| Beta suppression (%) | -32.34 ± 8.04 | -27.73 ± 8.60 | **< 0.001** |
| Latency (ms) | 305 ± 86 | 249 ± 82 | **< 0.001** |
| Beta rebound (%) | 55.35 ± 26.38 | 46.88 ± 24.31 | **< 0.001** |
| Latency (ms) | 756 ± 128 | 754 ± 129 | 0.95 |

VL, the vastus lateralis, VM, the vastus medialis.

### Effect of stimulation intensity on cortical and muscular response strengths

#### Muscular response to proprioceptive stimulation

Figure 2B and C show reflex force and peak-to-peak EMG amplitude of the vastus medialis and lateralis muscles in both intensity conditions. As expected, the peak EMG amplitude of VL muscle showed a statistically significant greater in higher intensity than in lower intensity condition (W = 54, p = 0.04), whereas there were no statistically significant differences in the peak EMG amplitude of VM muscle between both intensity conditions (W = 74, p = 0.15). Similarly, the reflex force was also significantly greater under higher intensity condition (W = 14, p < 0.001). Table 1 summaries the results for reflex force and peak EMG amplitude.

#### Evoked fields to proprioceptive stimulation

Figure 3 shows the evoked field results for both intensity conditions. The evoked activity appeared to peak over the contralateral leg representation area of the SM1 cortex in both intensity conditions, ∼58 ms after stimulus onset. MANOVA revealed a statistically significant main effect for intensity condition (F_4,37_ = 2.75, p = 0.04). Follow-up tests showed that peak evoked amplitude was significantly larger in higher intensity than in lower intensity condition (t = 2.52, p = 0.03), whereas the latency of peak evoked field did not differ significantly between both intensity conditions (W = 94, p = 0.53). Table 1 presents all evoked field results.

**Figure 3.**
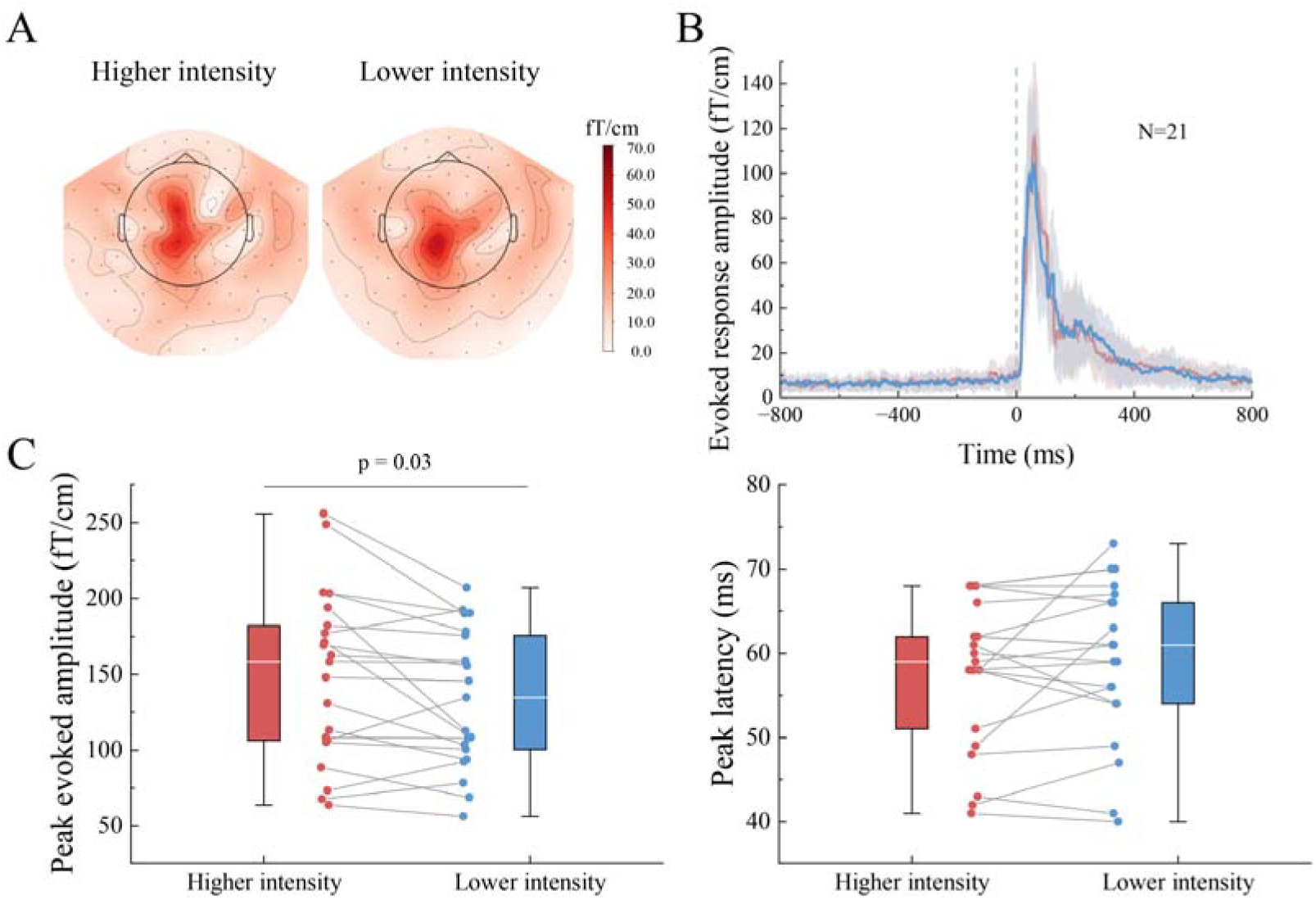
Cortical evoked responses to proprioceptive stimulation. (A) Group level evoked field topographies across participants are represented for higher and lower intensity conditions. (B) Grand average time-series of evoked fields for both intensity conditions. Vertical grey dashed line at time zero reflects stimulus onset. (C) Boxplots of peak values for peak evoked amplitude (left) and peak latency (right).

#### Induced response to proprioceptive stimulation

Figure 4 shows the induced response for both intensity conditions. Figure 4 left panels illustrate group-level spatial distribution for beta suppression and rebound for higher and lower intensity conditions. Surprisingly, the sensors of peak beta suppression were localized bilaterally more towards the hand area of the SM1 cortex at around 270 ms, after stimulus onset, whereas peak rebound activity was seen over the leg representation area at around 755 ms in both intensity conditions. Peak relative strength for beta suppression and rebound in the right panels were significantly stronger in higher intensity than lower intensity condition (t = −4.65, p < 0.001; W = 6, p < 0.001, respectively). However, the latency of the induced responses showed a different pattern: beta suppression peaked significantly earlier in lower intensity than in higher intensity condition (t = 5.69, p < 0.001), but no statistically significant differences were found in the latency of beta rebound between both intensity conditions (t = 0.067, p = 0.95). Table 1 shows all induced response results.

**Figure 4.**
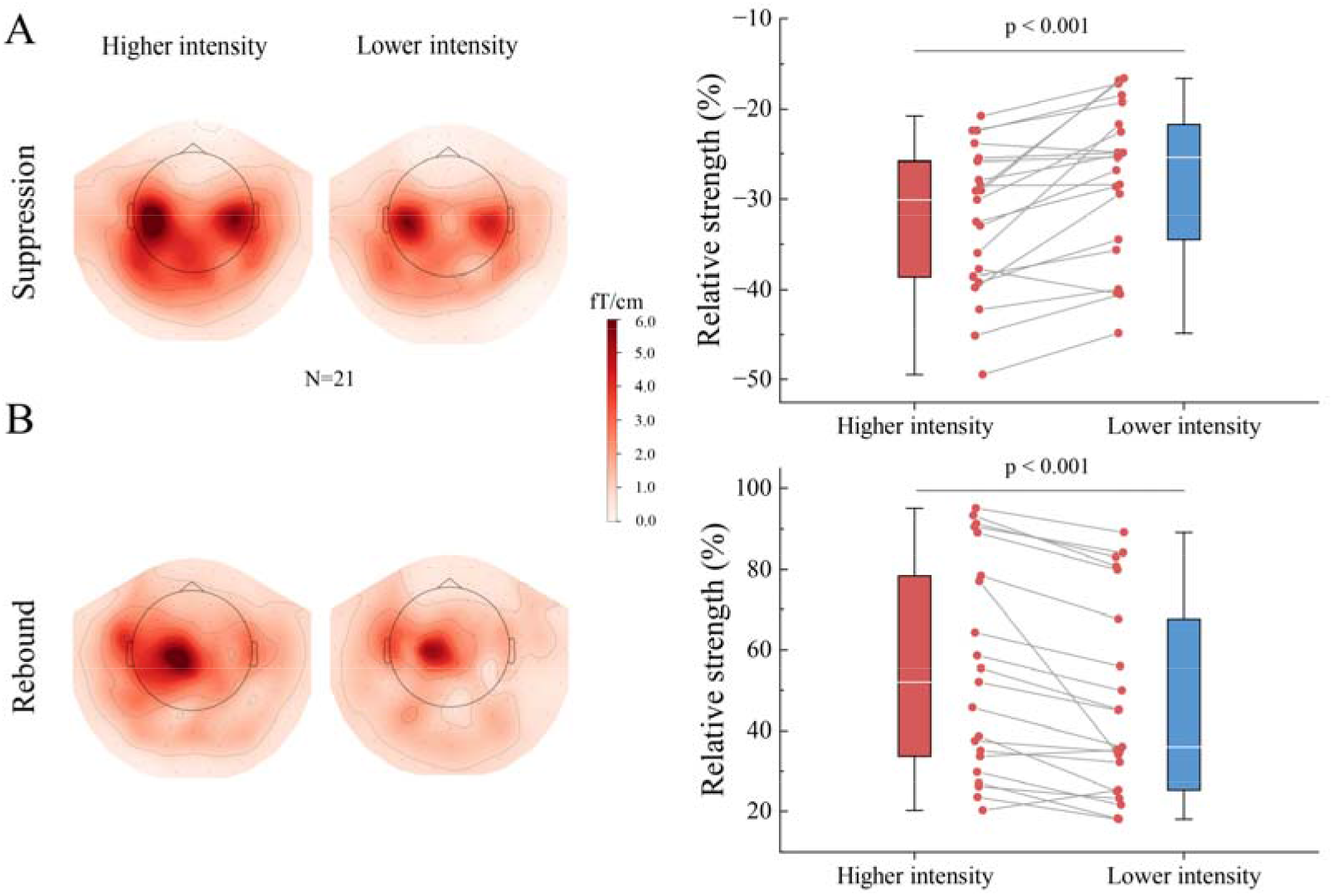
Induced cortical responses to proprioceptive stimulation in both higher and lower intensity conditions. (A) Topographies of group level induced response (left) and relative strength as boxplots (right) in beta suppression for higher and lower intensity conditions. (B) Topographic maps (left) and relative strength (right) as boxplots for beta rebound for both intensity conditions. Note that beta suppression is positive in the topography plots as the root-mean-square of the gradiometer pairs are plotted.

### Effects of stimulus intensity on response variability

Figure 5 shows muscular and cortical response variability for both intensity conditions. Figure 1D illustrates an example of averaged evoked field and muscular response based on 4 stimuli (i.e. one window) and all ∼100 stimuli during higher intensity condition. Notably, muscular response variability was significantly greater compared to evoked field variability in both intensity conditions (p < 0.001). However, there were no statistically significant differences in either evoked field (t = 0.76, p > 0.05) or in muscular response (F_2,39_ = 0.16, p = 0.85) between the stimulation intensities. Finally, the cortical and muscular response variabilities did not covary in either intensity conditions (p > 0.05, Fig. 5C). Table 2 summaries the response variability results.

**Figure 5.**
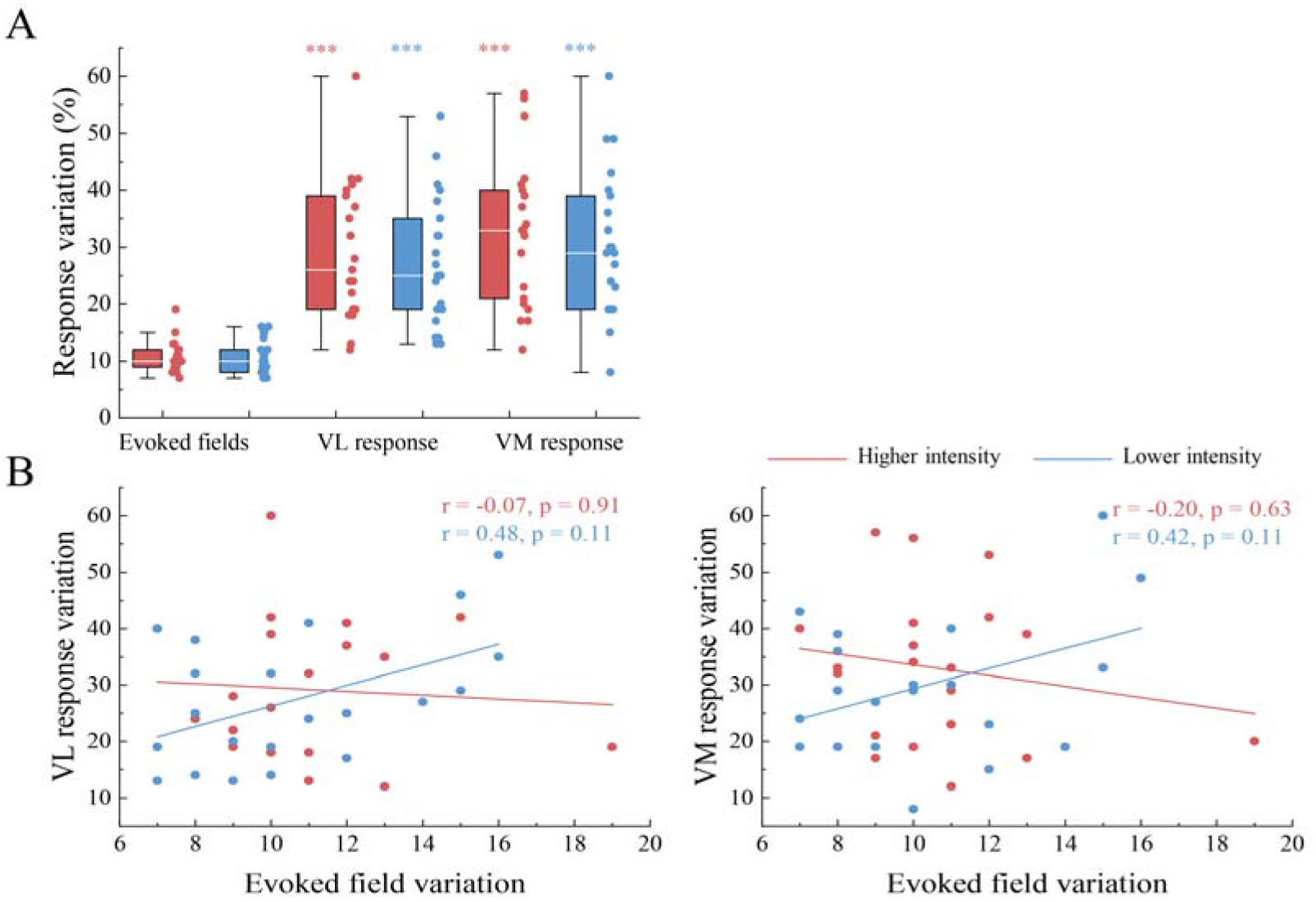
Response variability for cortical evoked fields and muscular response in higher and lower intensity conditions. (A) Sliding-window coefficient of variation (i.e., CoV value) as boxplots for evoked fields and VL, VM muscular responses across participants for both intensity conditions. Asterisks above muscular response variation bars indicate significant differences from evoked fields variation at the corresponding stimulus intensity. ***p < 0.001. (B) Spearman’s correlation coefficients between evoked fields and muscular response variability (VL muscle, left panel; VM muscle, right panel) in higher (red) and lower (blue) intensity conditions.

**Table 2.** Muscular and cortical response variability (mean ± SD) for higher and lower intensity conditions.

| **Coefficient of variation (CoV)** | **Higher intensity** | **Lower intensity** |
| --- | --- | --- |
| Evoked field (%) | 11 ± 2 | 11 ± 3 |
| VL EMG amplitude (%) | 29 ± 12 | 27 ± 12 |
| VM EMG amplitude (%) | 33 ± 13 | 31 ± 13 |
| Evoked field-VL EMG correlation | -0.07 | 0.48 |
| Evoked field-VM EMG correlation | -0.20 | 0.42 |

VL, the vastus lateralis, VM, the vastus medialis.

## Discussion

We examined whether the intensity of patellar tendon stimulation amplifies cortical proprioceptive processing in similar manner as is previously shown for the stretch-reflex muscular responses (Chandrasekhar et al., 2013; Tham et al., 2013; Tsuji et al., 2021). In line with our hypothesis, we observed that higher intensity tendon taps elicited stronger evoked and induced cortical and muscular responses. Moreover, muscular response variability was considerably higher than the cortical response variability, and the variability did not covary between the cortex and muscle. In addition, the respective response variabilities were not altered in response to the stimulus intensity. Together, these findings suggest that stronger proprioceptive stimulus intensifies the cortical proprioceptive processing, and that the early component of the proprioceptive afference is largely independently processed across cortical and spinal levels, which serve separate mechanistic purposes.

### Stimulus intensity modulates cortical and muscular response strength

#### Effect of stimulus intensity on muscular response

In agreement with our hypothesis and strong earlier evidence, peak EMG amplitude of VL muscle and reflex force were significantly higher during higher than lower intensity conditions. Comparable results have been obtained in the previous studies that investigated the effects of different types of tapping angles on reflex amplitude or EMG parameters (Chandrasekhar et al., 2013; Tham et al., 2013; Tsuji et al., 2021). The tendon reflex is known to be a stimulus-response relation in healthy subjects (Zhang et al., 1999), meaning that the larger the tapping force or angle within a given range, the stronger the reflex force and EMG amplitude. In the present study, higher stimulation intensity produced a significantly larger peak acceleration magnitude, indicating a stronger mechanical stimulus delivered to the tendon. Notably, no significant correlations were observed between stimulus kinematics and response strength in either muscular or cortical responses. Therefore, the increase in reflex force and VL amplitude support higher intensity stimulation generating a stronger effective reflex input.

Mechanistically, the stronger tendon tap generates greater/faster stretch of the quadriceps muscles, thereby increasing the dynamic activation of primary muscle spindles (Edin and Vallbo, 1990; Grill and Hallett, 1995; Matthews, 1963). Subsequently, this would increase Ia afferent discharge and, in turn, amplify the spinal reflex response. However, it should be noted that Ia afferent input was not recorded directly in this study and therefore we cannot determine how spindle firing was modulated by stimulus intensity in our design. In addition, the intensity-related effect was more evident in VL muscle than in VM muscle. The lack of stimulus intensity effects in VM reflex amplitudes could be a result of structural and reflex excitability differences compared to VL, and thus this observation should be interpreted cautiously (Doguet and Jubeau, 2014; Grob et al., 2016).

Importantly, the sensitivity of tendon reflexes to mechanical input may also have clinical implications. For example, higher tendon reflex torque has been reported in neurological populations, including multiple sclerosis patients, under comparable tapping force condition (Zhang et al., 2000). Such findings suggest that controlled tendon stimulation may complement conventional tendon-reflex examination by providing a more objective quantification of reflex gain or reflex sensitivity, particularly in conditions with altered reflex excitability.

#### Effect of stimulus intensity on evoked field

In our study, stronger peak amplitude of the evoked field (∼11%) was observed in higher intensity compared with lower intensity condition. The amplitude of evoked fields is known to depend on both the synchronicity of the postsynaptic activation and the size of activated neuronal population (David et al., 2006; Hari, 1990). Accordingly, the increase in the evoke field may reflect more synchronized and/or more extensive activation within the SM1 cortex neurons. The larger muscular responses observed under higher intensity condition further suggest that the stronger mechanical input generated a stronger and/or more synchronous proprioceptive afferent volley reaching the cortex, particularly through muscle afferents that contribute to early cortical response to proprioceptive stimulation (Drews et al., 1998; Mima et al., 1996). Thus, the quantitative increase in proprioceptive afference probably explains the stronger evoked fields for higher intensity than lower intensity conditions. Furthermore, the larger evoked fields under higher intensity condition may simply reflect the stronger afferent volley, both in terms of number of afferents activated and in number of each afferent firing, which would lead to more effective proprioceptive input to the sensorimotor cortex.

This finding is in accordance with previous studies showing increase in cortical responses with electrical stimulation intensity of quadriceps muscles and median nerve (Hewitt et al., 2022; Jousmaki and Forss, 1998; Smith et al., 2003; Torquati et al., 2002). However, this intensity-response relationship is unlikely to be strictly linear. These studies have also reported that cortical response can reach a plateau or even decrease when stimulus intensity exceeds a certain threshold. This may partly be due to gradual activation of different types of somatosensory afferents (e.g., nociceptors, etc.) that may modulate the cortical activation. The intensity-response relationship is out of the scope of the current study as only two stimulus intensities were used. However, nociceptors were likely not activated at least as strongly as the stimuli used were naturalistic and relatively light. Thus, future studies are needed to examine the linearity of the intensity-response curve of mechanical stimulation and compare it to peripheral electrical stimulation.

Nurmi et al. (2023) demonstrated that movement evoked fields strength was not influenced by movement range of proprioceptive stimulation of the finger. On contrary, here we observed that stronger tendon tap in lower limb leads to stronger evoked fields. These contradictory observations may reflect neurophysiological differences between upper and lower limb motor control, as well as differences in the need for proprioception. For example, fine-motor control is emphasized in the upper limb (e.g., hand movements), compared to gross locomotor control in the lower limbs (Staines et al., 1998). The contradictory findings may also be explained by methodological differences. Specifically, the smooth passive movement of index finger produced by Nurmi et al. (2023) did not evoke stretch-reflex in the muscles, whereas the patellar tendon tap did. Thus, the cortical response to tendon tap reflects not only the initial proprioceptive afferent volley generated by the tap itself, but also the subsequent stretch-reflex evoked muscle “twitch” activating the proprioceptive afferents again about 28 ms after the initial external tap stimulus.

Beyond the evidence from evoked fields, CKC studies provide converging evidence that cortical proprioceptive responses depend on how peripheral input is generated and distributed. For example, simultaneous stimulation of multiple fingers enhances CKC strength compared with single finger stimulation, suggesting that more comprehensive proprioceptive afference can strengthen cortical coupling (Hakonen et al., 2022). Similarly, lower-limb CKC strength is modulated by passive movement (including hip, knee and ankle joint movements) parameters, especially in movement range and frequency (Zhao et al., 2025). Together, these findings suggest that the strength of cortical responses to proprioceptive stimulation may depend not only on the degree of peripheral afference, but also on the stimulation modality, its mechanical features and the joint/limb stimulated.

#### Effect of stimulus intensity on induced response

Beta suppression is typically associated with cortical activation and increased excitability (Hall et al., 2011; Neuper et al., 2006), whereas beta rebound is often interpreted as a post-event re-synchronization linked to reduced excitability or inhibitory processing in response to unvoluntary movement (Gaetz et al., 2011; Neuper and Pfurtscheller, 2001; Zhang et al., 2008). In the present study, higher intensity stimulation elicited stronger beta modulation than lower intensity stimulation, with a broader spatial distribution of the induced response, suggesting that higher intensity proprioceptive stimulation enhanced both cortical processing of proprioceptive afference and the subsequent reorganization of sensorimotor network activity. A possible explanation for stronger induced responses is that the stronger tendon tap generated a larger peripheral proprioceptive afference, consistent with the larger spinal stretch-reflex EMG amplitude observed under high intensity stimulation. Although muscular and cortical responses are processed at different levels, the larger EMG response indicates that higher intensity stimulation produced greater muscular stretch-reflex and thus likely a stronger proprioceptive afferent volley from the periphery towards the central nervous system. In turn, this stronger proprioceptive afference may have resulted in stronger recruitment of neuronal networks in the SM1 cortex. In addition, multimodal somatosensory afference may strengthen the beta suppression, since volitional finger extension and median nerve electrical stimulation have shown to produce stronger beta suppression than cutaneous tactile stimulation alone in index finger (Alegre et al., 2002; Houdayer et al., 2006). Furthermore, we cannot rule out the possibility that stronger intensity stimulation activated cutaneous receptors more effectively, as they will be inevitably activated during the tendon tap. The lower intensity stimulation was also likely strong enough to activate majority of the cutaneous receptors in the skin on top of the patellar tendon.

An EEG study using peripheral wrist extensors electrical stimulation by Insausti-Delgado et al. (2020) showed stronger beta suppression with an increment of the stimulation intensity over the SM1 cortex, being consistent with our observation; however, they did not observe a significant effect on beta rebound amplitude. In contrast, an increase in the movement range of passive finger movement stimulus did not alter MEG beta suppression and rebound strength (Nurmi et al., 2023). Together, these studies suggest that beta modulation may not only reflect an increase in stimulation magnitude but may instead depend on the nature of the elicited afference, shaped by different somatosensory stimulations.

It is suggested that beta suppression and rebound likely track partly distinct aspects of the cortical proprioceptive processing, since beta suppression and rebound have been shown to arise from functionally distinct neuronal populations (Parkkonen et al., 2015). This partial difference was reflected descriptively in the sensor-level topographical representations observed in the present study. The beta rebound peaked over the expected leg area of the SM1 cortex whereas peak beta suppression was located more laterally. A similar pattern was observed in our previous study (Li et al., 2026), in which additional source-level analyses suggested that beta suppression to proprioceptive stimulation in the knee joint may involve cortical regions beyond the classical SM1 leg region.

### Cortical and muscular response variability do not covary from stimulus-to-stimulus

Although stimulus intensity clearly amplified cortical and muscular responses, the intensity did not significantly influence their level of variability. This observation suggests that increasing the tendon-tap intensity amplified the response gain but did not alter the stimulus-to-stimulus stability of the underlying input processing at cortical and spinal levels.

At the group level, there was pronounced variability in the muscular EMG reflex response amplitudes with ∼30% CoV, but the cortical MEG responses showed markedly lower variability with ∼10% CoV in both intensity conditions. Interestingly, the cortical MEG and spinal muscular response variability did not covary, i.e. there was no statistically significant correlation between cortical and muscular responses variation value in either intensity conditions. Thus, the higher variability of muscular output was not mirrored in the cortical ones, suggesting that the early proprioceptive processing is highly distinct at spinal and cortical levels.

This pattern may reflect both pathway-level and functional differences between spinal and cortical processing of proprioceptive afference. The patellar tendon reflex provides a rapid and adaptable spinal output in which muscle spindle afferent input activates the motoneuron pool and generates the quadriceps response. This output is likely to be sensitive to trial-to-trial fluctuations in tendon-tap delivery, muscle-tendon mechanics, spindle sensitivity, fusimotor control and motoneuronal recruitment (Blum et al., 2020; Dimitriou, 2022; Macefield and Knellwolf, 2018). However, the cortical evoked fields reflect proprioceptive afference after additional processing along the dorsal column pathway, including relays in the medullary nucleus gracilis and the ventral posterolateral thalamus before reaching the SM1 cortex (Al-Chalabi et al., 2026; Navarro-Orozco et al., 2026). These ascending relays and intracortical integration stages may mean that local peripheral or reflex-related fluctuations are not directly expressed in the population-level cortical response.

Functionally, this distinction is consistent with the different roles of spinal and cortical proprioceptive processing. The spinal cord maintains a flexible and quickly adjustable reflex gain to protect joint function under varying conditions, whereas the cortex needs accurate feedback about the current state of the locomotor system to plan and learn motor actions. (Proske and Gandevia, 2012; Scott, 2004). This division of function highlights not only a pathway-level difference in how the afferent signal is transformed before reaching the cortex, but also how local adaptability at the spinal level works together with more consistent processing in the cortex, allowing the sensorimotor system to balance immediate reflex needs with stable higher-order control.

Finally, these findings indicate that response variability was shaped more by the physiological level of processing than by stimulus intensity itself. Although cortical and muscular responses were initially elicited by the same tendon-tap evoked proprioceptive afference, the larger variability observed in muscular response was not mirrored in cortical evoked fields. With the non-significant correlation results, this pattern supports the functional dissociation between spinal and cortical processing of proprioceptive afference.

### Further perspectives and limitations

In the current stimulus-to-stimulus variability analysis, a sliding-window CoV method had to be used since cortical evoked fields have limited signal-to-noise ratio in MEG due to massive “brain noise” from activity not related to the task. Therefore, the CoV value should be interpreted as a window-estimated response variability with four stimuli, rather than as direct single stimulus variability. With this approach, some stimulus-to-stimulus fluctuations are lost in the averaging, and the actual variation is likely larger. Nevertheless, the robust estimation of MEG evoked response with only four stimuli is remarkable, emphasizing the strong activation of the SM1 cortex neurons to proprioceptive stimulus, and suggesting that this method is feasible to be used in future studies.

The present study tested only two stimulation intensities and did not systematically characterize the full stimulus-response relationship. As a result, the current data showed that stronger tendon taps increased the response strength without changing response variability, but did not establish whether this pattern would remain, plateau or reverse at higher or more finely graded intensities. This is particularly relevant because the reflex output to patellar tendon may not be further increased beyond a certain stimulation range. Additionally, a recent proprioceptive study suggested that some cortical measures can remain stable despite variation in movement parameters (Nurmi et al., 2023). Future work with a more finely graded intensity setup would allow a more mechanistic account of how peripheral input strength is transformed across spinal and cortical levels.

Although higher and lower intensity conditions were separable within individuals, peak acceleration magnitude of the tendon tap still varied across participants. The between-subject variability is difficult to eliminate fully for the patellar tendon reflex. Participants differ in body height, leg length, local tendon anatomy and geometry, and their most sensitive tapping point for evoking the stretch reflex. Consequently, even with standardized stimulation settings, the effective mechanical input may not be strictly identical across individuals. This variability may influence the absolute stimulus level of inter-individual intensity with different sensitivity thresholds across participants, but it does not have the effect on within-subject intensity. For this reason, the stimulus intensity in the current study should not be understood strictly in absolute terms, but as more relative terms as difference between lower and higher intensity conditions. To improve towards more constant stimulus intensity across participants in the knee-joint stimulator, more sophisticated adjustments would be required for the MEG chair and the MEG-compatible stimulator, including more individualized and fixed adjustment of leg position and hammer trajectory, or completely new pneumatic cylinder-based stimulator, for example.

Finally, our findings facilitate the understanding how proprioceptive afference is processed across different levels of the proprioceptive afferent pathways. Cortical proprioceptive processing has already been shown to alter with ageing (Piitulainen et al., 2018; Walker et al., 2020), Parkinson’s disease (Vinding et al., 2019), cerebral palsy (Illman et al., 2024) and type I diabetes mellitus (Mujunen et al., 2025). However, it is unclear how response variability to proprioceptive stimulation at cortical and spinal levels alters in various motor disorders. This may help clarify whether impaired proprioceptive function in ageing or disease primarily reflects altered peripheral encoding or disrupted cortical integration.

## Conclusion

Higher intensity proprioceptive stimulation of the patellar tendon elicited stronger cortical and muscular responses when compared to lower intensity condition, suggesting that the respective cortical processing of proprioceptive afference is intensified with stimulus intensity. As expected, the cortical evoked field peaked in the contralateral leg region of the SM1 cortex, whereas, unexpectedly, the beta suppression, reflecting activation of the cortex, peaked bilaterally towards the contralateral and ipsilateral hand regions in both intensity conditions, revealing wider neuronal network processing knee proprioception than expected in the SM1 cortex. The stimulus intensity did not affect the stimulus-to-stimulus cortical or muscular response variability, but the variation value was strikingly stronger in the spinal level muscular responses than the cortical level evoked fields, suggesting that the cortical and spinal proprioceptive processing are largely distinct. It appears that the cortex needs intact proprioceptive feedback for sensorimotor integration, whereas spinal level response is susceptible to complex neuronal modulation crucial for finetuning the motor plane before the final common pathway to the motor units of the muscle.

## Acknowledgements

The authors gratefully acknowledge Markus Kerminen for his precious technical assistance in building the device at the University of Jyväskylä, and Heidi Pesonen, Hanne Ahonen for the valuable assistances with data collection.

## Funding Statement

This work was supported by Academy of Finland grant (grant #361732) to HP. This work was also supported by the PhD scholarship from China Scholarship Council (#202306060029) to FL.

## CrediT authorship contribution statement

**Feiyue Li:** Writing – review & editing, Writing – original draft, Visualization, Formal analysis, Data curation, Software. **Anni Byman:** Writing – review & editing, Validation, Methodology, Investigation, Data curation. **Junru Chen:** Writing – review & editing, Data curation. **Toni Mujunen:** Writing – review & editing, Validation, Methodology, Investigation, Data curation, Software. **Harri Piitulainen:** Writing – review & editing, Supervision, Resources, Project administration, Methodology, Funding acquisition, Conceptualization.

## Declaration of competing interest

The authors declare that they have no known competing financial interests or personal relationships that could have appeared to influence the work reported in this paper.

## Abbreviations

SM1: primary sensorimotor cortex
MRI: magnetic resonance imaging
EEG: electroencephalography
MEG: magnetoencephalography
CKC: corticokinematic coherence
ERD: event-related desynchronization
ERS: event-related synchronization
EMG: electromyography
VL: vastus lateralis
VM: vastus medialis
TSE: temporal spectral evolution
CoV: coefficient of variation
RMS: root mean square.

## Notes

### Competing Interest Statement

The authors have declared no competing interest.

